# Development of an on-chip detection of Zika virus and antibodies simultaneously using array of nanowells

**DOI:** 10.1101/302893

**Authors:** Touyana Semenova, Alexandria Voigt, William Donelan, Alek Aranyos, Janet Yamamoto, Mobeen R. Rathore, Cuong Q. Nguyen

**Affiliations:** Department of Infectious Diseases and Immunology, College of Veterinary Medicine, University of Florida, Gainesville Florida, USA.; University of Florida Center for HIV/AIDS Research, Education and Service (UF CARES), University of Florida, Gainesville Florida, USA.; Department of Oral Biology, College of Dentistry, University of Florida, Gainesville Florida, USA.; Center of Orphaned Autoimmune Diseases, University of Florida, Gainesville Florida, USA.

**Keywords:** Zika virus, microengraving, real-time PCR, on-chip nanowells, diagnoses, serology

## Abstract

Zika virus (ZIKV) infections are an emerging health pandemic of significant medical importance. ZIKV appeared recently in the Americas from Africa via the South Pacific. The current outbreak has garnered attention by exhibiting unique characteristics of devastating neurodevelopmental defects in newborns of infected pregnant women. Current guidelines for ZIKV diagnostics developed by the Center of Diseases Control and Prevention (CDC) consist of nucleic acid testing, plaque reduction neutralization test (PRNT), and a serologic test for IgM detection. To better accommodate and comply with these guidelines, we developed a simultaneous on-chip detection of ZIKV and anti-ZIKV antibodies using an array of nanowells. Using on-chip microengraving, we were able to detect anti-ZIKV antibodies and their immunoglobulin isotypes. In parallel, applying on-chip real-time PCR with epifluorescence microscopy, we were able to quantify ZIKV viral load as low as one copy. To test clinical samples of patients at the post-convalescent stage, we analyzed samples from 8 patients. The on-chip nanowells could effectively identify antibodies that reacted against ZIKV envelope protein and their isotypes with high sensitivity and specificity. The small sample requirement with high specificity and sensitivity and combined molecular and serological tests could potentially be very advantageous and beneficial in accurate detection of Zika infection for better disease monitoring and management.

## INTRODUCTION

After the first reported human infection outbreak on Yap island in 2007, Zika virus (ZIKV) spread dramatically in the Pacific Ocean by a larger epidemic in French Polynesia in 2013-2014 with 32,000 estimated infections (1, 2) and subsequent outbreaks on other Pacific Islands and in the Americas (1, 3–7). Ninety-five countries have been classified by the CDC as risk areas for ZIKV transmission, and 47 countries and territories in the Americas reported ZIKV outbreaks during 2015-2016 (3, 8). In recent years, ZIKV became a serious cause for public health due to its teratogenic and neuropathic outcome in infants and neurological disorders such as Guillain-Barré syndrome in adults (1, 4, 7, 9–11).

ZIKV is a single-stranded RNA arbovirus (Family *Flaviviridae*, genus *Flavivirus*) transmitted by Aedes mosquitos. The ZIKV genome contains 10,941 nucleotides encoding 3,419 amino acids with 5’ and 3’ non-coding regions (NCR) and one open reading frame. The open reading frame encodes a single polyprotein that is later cleaved into three structural components: capsid (C), precursor membrane (prM), and envelope protein (E); and seven nonstructural (NS) proteins: NS1, NS2A, NS2B, NS3, NS4A, NS4B, and NS5 (12). Hierarchical cluster analysis shows that ZIKV and Dengue virus (DENV) cluster at a higher hierarchical level and that ZIKV is phylogenetically most related to Spondweni virus (12, 13). This separation of the ZIKV cluster from other flavivirus clusters at a similar hierarchical level may play an important role on pathogenesis and tissue tropism despite the similarity of clinical symptoms to other flavivirus infections (13). Contrary to other flaviviruses, ZIKV can be transmitted vertically by sexual contact and intravenous transfusion. Consequently the virus can be present in human aqueous (14), seminal fluid (14, 15), urine (15), vaginal secretions (16), breast milk, amniotic fluid (5, 17), fetal cerebrospinal fluid, cord blood, infant blood at the second day of birth, and placenta (17). Additionally, ZIKV infection has recently been described in posttransplant patients of solid organs and stem cells from asymptomatic infections to meningoencephalitis (18).

Current guidelines of ZIKV diagnostics developed by the Center of Diseases Control and Prevention (CDC) consist of testing for ZIKV antibodies using the IgM antibody capture enzyme-linked immunosorbent assay (MAC-ELISA) followed by validation using plaque reduction neutralization testing (PRNT). Additionally, nucleic acid testing (NAT) should be performed during the first 6 weeks after the onset of symptoms (19). However, final results may be misinterpreted due to ZIKV epidemiological characteristics and diagnostic limitations. In generally ZIKV is characterized by having an asymptomatic course (20, 21), short transient incubation, and viremic periods (3-14 days, median: 6.2 days and 5 days respectively) (22, 23). Viremia may fluctuate depending on samples tested (whole blood, serum, urine, semen, or amniotic fluid) (21, 24). Furthermore, detection of viral RNA can be prolonged in pregnant women and in adults with Guillain-Barré syndrome (25–29). Laboratory results for serological testing of IgM against ZIKV may sometimes be difficult to interpret, especially for pregnant women, due to possible long persistence of IgM against ZIKV (2-4 months) (19, 30). For competent management of infection, the CDC has recommended that it is necessary to concurrently obtain a patient-matched serum specimen for NAT and/or IgM serological tests. Therefore, in this study we proposed the development of a novel diagnostic method based on massively parallel on-chip detection of ZIKV using a modified fluorescent polymerase chain reaction (PCR) and isotypic anti-ZIKV antibodies by microengraving in nanowells. The results indicated that this on-chip molecular biology test exhibited significant sensitivity for detection of low viral copy number. Simultaneously, the microengraving serological test was able to identify anti-ZIKV antibodies and their isotypes. Therefore, utilization of on-chip detection using nanowells might provide a significant technological advantage which benefits the monitoring and clinical management of ZIKV infection.

## MATERIAL AND METHODS

### Patient materials

Serum and plasma samples were purchased from Boca Biolistics (Pompano Beach, USA). These clinical samples were tested for ZIKV RNA by real-time PCR (COBAS Z480 PCR instrument and Light Mix Modular Zika Virus PCR Real Time, Roche, Switzerland) and for IgG/IgM antibody reactive with ZIKV antigens by Euroimmun IgG and IgM EIA (Euroimmun AG, Germany). Furthermore, samples were tested for West Nile virus (WNV), Chikungunya virus (CHIKV), and Dengue virus (DENV) RNAs and virus-specific antibodies by real-time PCR and ELISA according to the manufacturer’s instructions (InBios International, Inc., Seattle, WA). All procedures were reviewed and approved by the University of Florida Institutional Review Board.

### Fabrication of arrays of nanowells

Sylgard 184 silicone elastomer base (polydimethyl-siloxane, PDMS) and curing agent with a 10:1 weight ratio was combined and mixed vigorously. The mixture was degassed under vacuum for 2 hrs and poured into a custom-built aluminum mold containing a silicon wafer with a patterned array of posts. The mixture was set to cure for 2 hrs at 80°C and adhered directly to a 3”×1” glass slide. The pattern on the master aluminum mold was transferred to the cured PDMS in bas-relief. In this experiment, a master aluminum mold was used that contained blocks of 7×7 nanowells, 4×4 blocks and 6 columns×18 rows with nanowell dimensions of 50 μm×50 μm×50 μm for a total of 84,672 nanowells per array.

### Real-time PCR amplification of ZIKV and DENV

ZIKV RNA was extracted from 140 μl of serum and plasma samples using the Qiamp Viral RNA Mini RNA kit (Qiagen, Hilden, Germany) according to the manufacturer’s instructions. Primers (Forward: 5’d CAGCTGGCATCATGAAGAAYC 3’; Reverse 1: 5’d CACTTGTCCCATCTTCTTCTCC 3’; Reverse 2: 5’d CACCTGTCCCATCTTTTTCTCC 3’) and probe (5’d FAM-CYGTTGTGGATGGAATAGTGG-BHQ-1 3’) were designed according to the previous study (31) and purchased from LGC Biosearch Technologies (Petaluma, CA). A mastermix contained: 5 μl iTaq universal probes reaction mix, 0.25 μl of iScript reverse transcriptase, 100 nM for each primers, 150 nM for each probes, 2.75 μl of nuclease-free water and 1 ng viral RNA. Mastermix was deposited on a 96 well PCR plate and sealed with PCR plate sealing film. Real-time PCR was performed on the CFX96 Touch real-time PCR detection system (Bio-Rad, CA). A thermal cycling protocol was as followed: reverse transcription at 50 °C for 10 min, polymerase activation and DNA denaturation at 95 °C for 3 min followed by 40 cycles of amplification: denaturation at 95 °C for 15 sec, annealing/extension with plate reading at 60 °C for 30 sec. Similar protocols were used for DENV RNA detection using the FDA-approved CDC DENV-1-4 RT-PCR assay (32) performed on the CFX96 Touch real-time PCR detection system (Bio-Rad, CA).

### Plaque reduction neutralization test

The presence of neutralizing antibodies was determined as previously described (33). ZIKV Puerto Rico strain PRVABC59 was used for this assay. Patients’ samples, positive, and negative controls were titrated with media containing Eagle’s Minimum Essential Medium (EMEM) (Corning, NY), 2.5 % fetal bovine serum (FBS, Atlanta Biologicals, GA), and gentamicin at 25μg/mL (Gybco,BRL, NY), and viral stock was added into a 96 well plate and incubated for 1 hour at 37ºC with 5% CO_2_. Vero cells (ATCC CCL 81, epithelial cells of African green monkey—*Cercopithecus aethiops*; Manassas, VA) were plated in a 96 well plate at 90% monolayer confluency. The growth media was removed from the 96 well plate and patients’ samples with viral stock were deposited into the plate with Vero cells. After 48 hours of incubation at 37ºC with 5% CO_2_, unabsorbed virus was removed, and methylcellulose overlay medium containing EMEM, 7% NAHCO_3_, and gentamicin (25 μg/mL) was added to each well. After 48 hours of incubation at 37ºC with 5% CO_2_, the plate was stained by crystal violet solution containing crystal violet, methanol, and distilled water, and the plaques were counted. According to the CDC guidelines, (19) a titer <10 is considered as negative, and a titer ≥ 10 is considered positive.

#### On-chip microengraving

Capture slides were coated for 1 hour with anti-mouse Abs (for mouse monoclonal antibodies: anti-ZIKV E clone ZV-2 and anti-flavivirus clone D1-4G2-4-15) or anti-human Abs (human sera and plasmas) as previously described(34, 35). Anti-ZIKV envelope protein ZV-2 was added to the nanowells (NR-50414, BEI Resources) as positive control and anti-flavivirus clone D1-4G2-4-15 served as negative control (NR-50327, BEI Resources). After incubation capture slide was blocked in 3% milk buffer solution for 1 hour, rinsed with PBST, PBS, deionized water, spun dry, and stored at 4°C (36). Ten microliters of serum or plasma sample was deposited onto the nanowells and hybridized with the treated capture slide for 2 hours in a hybridization chamber (Agilent Technologies, CA) at RT. After incubation, the capture slide was processed using Tecan Pro HS 4800 Hybridization Station (Tecan, Männedorf, Switzerland) by adding a mixture of goat anti-mouse IgM-PE, anti-mouse IgG-Alexa Fluor (AF)647, anti-mouse IgA-AF555 (mouse monoclonal antibodies) or goat anti-human IgM-PE, anti-human IgG-AF647, anti-human IgA-AF555 (human sera/plasmas) (SouthernBiotech, AL), and ZIKV envelope protein ectodomain (Protein Science Corporation, CT) conjugated with AF488 using DyLight Antibody Labeling kit (Thermo Scientific, IL). Capture slide was scanned using the Genepix 4400A scanner (Molecular Devices, CA).

#### On-chip real-time PCR

A mastermix was prepared containing 5 μl iTaq universal probes reaction mix, 0.25 μl of iScript reverse transcriptase, 100 nM for each primers, and 150 nM for each probes as presented previously, 2.75 μl of nuclease-free water, and 1 μl serum/plasma. Zika RNA or Zika virus were used as positive controls, and healthy donor sera or no template were used as negative controls. The mastermix was deposited on the nanowell chip. The chip was sealed with the Frame-Seal^™^ in Situ PCR and Hybridization Slide Chambers and placed on the Eppendorf^™^ In Situ Block Adapter for Mastercycler^™^ Thermal Cycler (Eppendorf, Hamburg, Germany) to run one-step realtime-PCR. A thermal cycling protocol was as followed: reverse transcription at 50 °C for 10 min, polymerase activation and DNA denaturation at 95 °C for 3 min followed by 40 cycles of amplification: denaturation at 95 °C for 15 sec, annealing/extension with plate reading at 60 °C for 30 sec. After PCR, the microarray chip was analyzed for detection of signal and quantification of fluorescent intensity using an automated epifluorescence microscope equipped with a phase contrast, motorized stage, 405-nm and 488nm wavelength filter sets using Nikon NIS-Elements Advanced Research image capture software (Nikon, NY).

### Data and statistical analyses

The mean fluorescent intensity (MFI) for each well with a positive signal were generated using GenePix Pro7 Software (Molecular Devices, CA). NIS-Elements Microscope Imaging Software (Nikon, NY) was used to quantify the MFI of real-time PCR results. Data was analyzed using the unpaired two-tailed Mann-Whitney test (GraphPad Prism, CA) to determine the statistical significance. In all cases, p values ≤ 0.05 were considered significant. Excel (Microsoft, WA) was used to perform regression analysis.

## RESULTS

### Microengraving for anti-ZIKV antibodies using the on-chip nanowells

Diagnosis of ZIKV is typically based on nucleic acid amplification to enumerate the viral load or immunoassays to determine the antibody response to the virus. Nucleic acid amplification and antibody determination are routinely performed separately, since there is no technique that detects both parameters simultaneously. Therefore, we sought to determine if we could perform ZIKV amplification and detect ZIKV-specific Abs using the on-chip nanowells. As a proof-of-concept, we utilized monoclonal anti-ZIKV clone ZV-2, which has been shown to recognize ZIKV envelope protein (ZIKV E) as a positive control and monoclonal anti-flavivirus clone D1-4G2-4-15 as negative control, which has neutralizing ability against ZIKV, but does not bind to the viral envelope protein. Abs were serially diluted at 1:10, 1:50, and 1:100 and 10 μl of undiluted and serially diluted Abs were deposited onto the nanowell chip and hybridized with a capture slide coated with goat anti-human Ig and goat anti-human IgG (H+L). Detection antibodies conjugated with specific fluorochromes were added to the capture slide microarray. As presented in Figure 1A, monoclonal anti-ZIKV clone ZV-2 was able to bind to ZIKV E protein as anticipated. Additionally, the microengraving process was able to detect the IgG and IgA, but not IgM isotypes. Monoclonal anti-flavivirus clone D1-4G2-4-15 which has been shown to not react against the E protein, was negative for E protein, IgA, and IgM, but positive for IgG isotype using on-chip microengraving. Regression analysis showed that anti-ZIKV E dilution was positively correlated with fluorescent intensity (R^2^=0.7909) (Figure 1B). Therefore, the data indicated on-chip microengraving can be utilized to concomitantly detect ZIKV-specific antibodies and the isotypes present in the sample.

**Figure 1.**
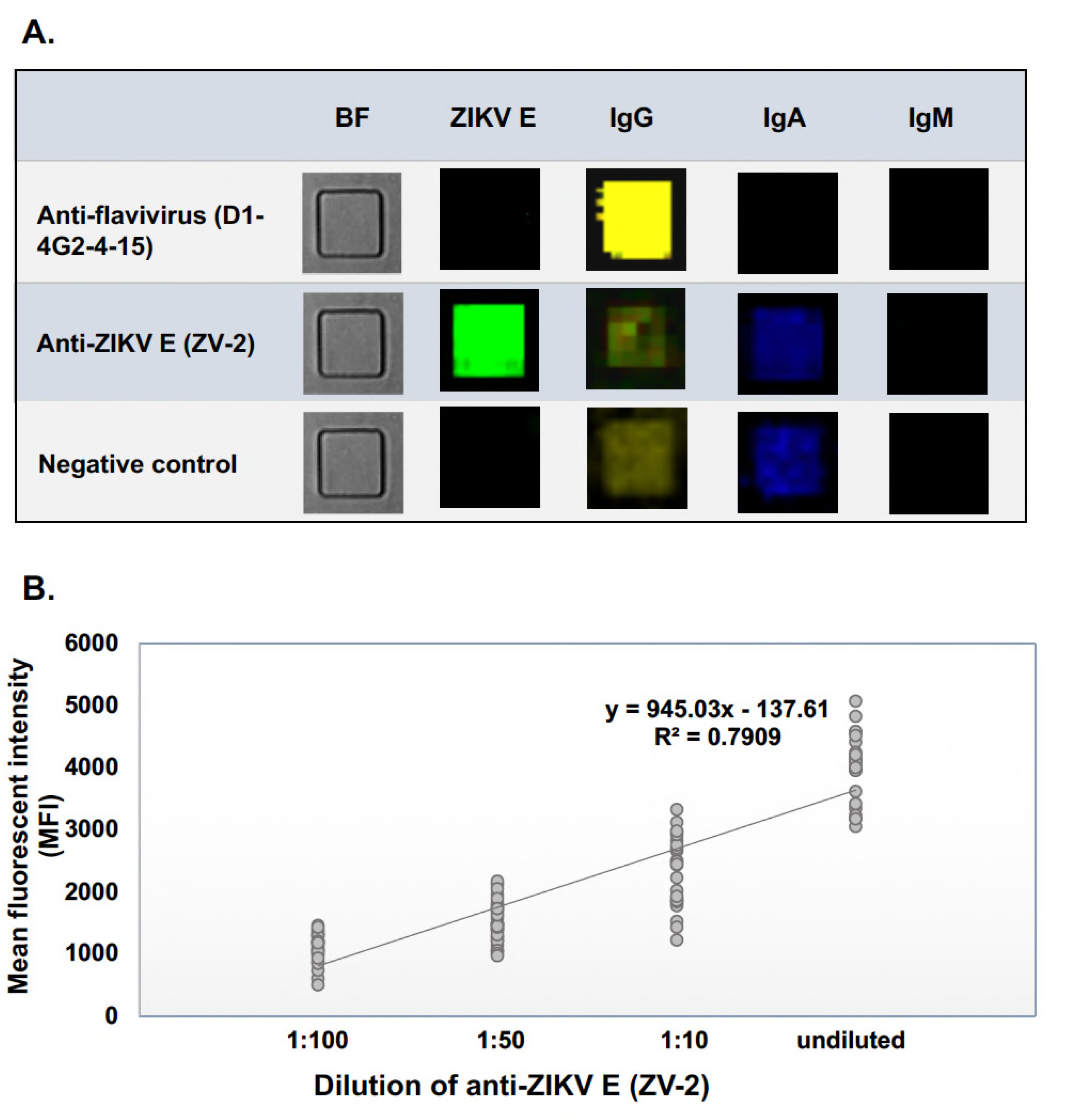
Detection of anti-ZIKV using the on-chip microengraving process. **A.** Samples were deposited in the nanowells. Microengraving was used to determine the reactivity for ZIKV E protein and Ig isotypes using fluorochrome-labeled proteins (ZIKV E: AF488, IgM: PE, IgG: AF488, and IgA: AF555). Representative microarray micrograph of capture slides showed the reactivity of the positive control (monoclonal anti-ZIKV clone ZV-2) and negative control (monoclonal anti-flavivirus antigen clone D1-4G2-4-15 and detection secondary proteins alone). Healthy human donor sera were also used as negative control for ZIKV E protein. **B.** Monoclonal anti-ZIKV clone ZV-2 was diluted at 1/10, 1/50, and 1/100 and subjected to microengraving. MFI for each nanowells at each dilution were determined using GenePix Pro7 Software and presented. The experiments were performed at least five times for consistency.

### Real-time PCR for ZIKV using the nanowell chip

Significant progress has been made in molecular detection of viruses. The optimization of detection methods using real-time PCR based assays allows for assays to be performed rapidly and produce specific, sensitive, and reproducible results for virus detection. However, the real-time PCR based assays still have limitations, particularly the sample volume and threshold of detection. To address these specific challenges, we performed the real-time PCR assay on the nanowell chip with a limited number of viral copies and volume. Undiluted ZIKV samples were serially diluted in plasma of healthy control and different viral copy numbers (1, 10, 100, or 1000) were deposited into the each individual nanowell predicted by the Poisson distribution. A mastermix containing the reverse transcriptase, polymerase, primers, and probes specific to ZIKV were added to the nanowell chip and placed on a standard laboratory PCR instrument. Using epifluorescence microscopy, the chip was imaged to examine the change in fluorescent signals based on the change in viral loads (Figure 2A). As indicated in Figure 2B, the nanowell chip was able to capture fluorescent intensity from 1 copy to 1000 copies and correlated strongly by linear regression analysis (R^2^=0.9631). DENV was not detected using ZIKV primer sets (data not shown). The result demonstrates the sensitivity and specificity of the nanowell chip for ZIKV detection, and it can be used to quantify the exact viral load based on fluorescent intensity.

**Figure 2.**
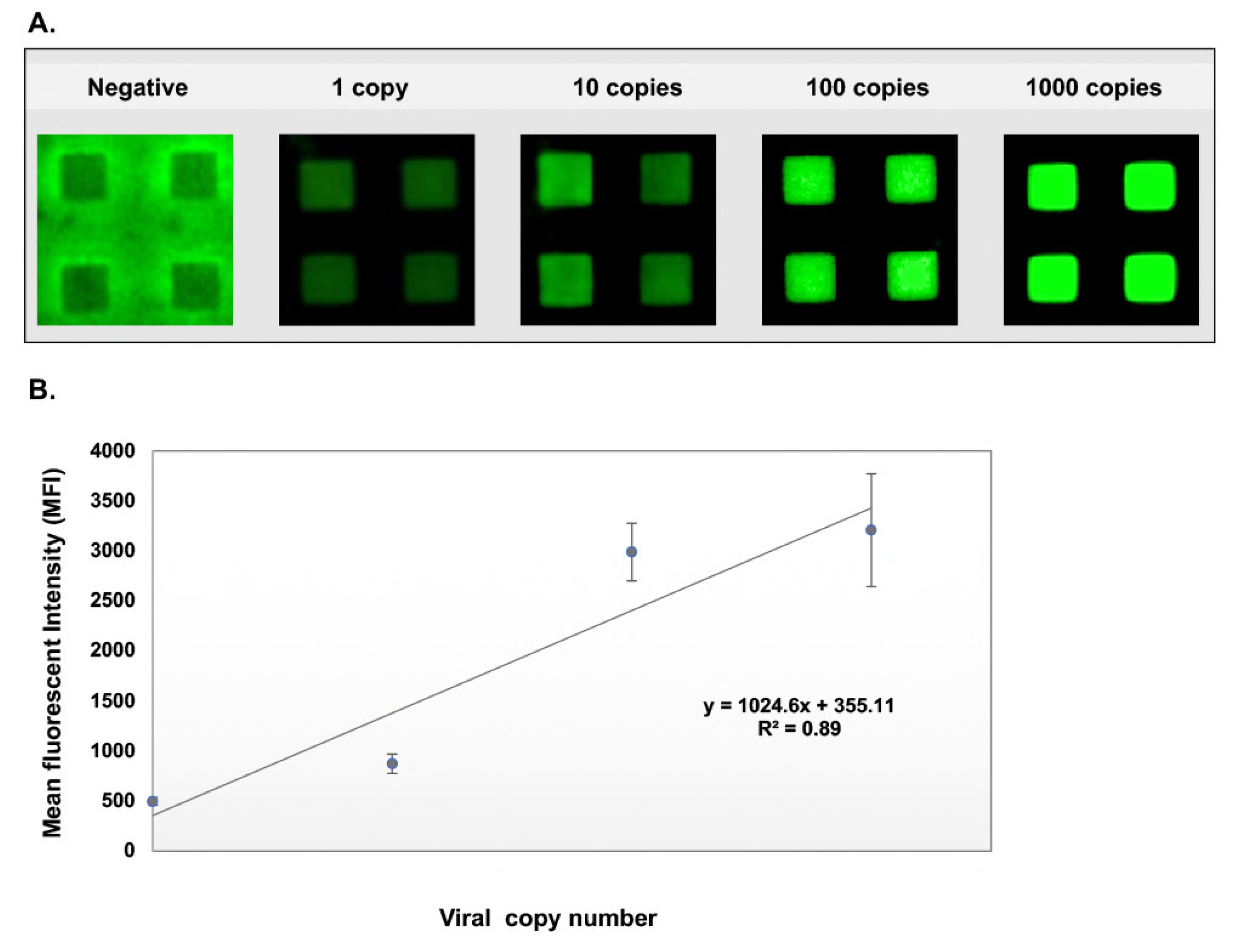
Detection of ZIKV using on-chip real-time PCR. **A.** Healthy human donor sera (n=5) were used. The actual number of viral copies that deposited in each nanowell can not be controlled, but the proportion of nanowells deposited with different numbers of viral copies of starting viral load can be predicted using the Poisson distribution. The optimal viral copy concentration and real-time PCR reagent volume were determined empirically to obtain the highest percentage of nanowells containing different copy numbers of ZIKV (1, 10, 100, and 1000 copies). Samples were deposited in the nanowells and a mixture of primers, probe, and PCR reagents were added. PCR was performed using a customized Eppendorf thermal cycler and nanowells were imaged using an automated epifluorescence microscope. Representative images of nanowells at different dilutions of virus is presented. **B.** MFIs of each nanowell on the chip were analyzed by Nikon NIS-Elements Advanced Research image capture software (Nikon, NY) and presented as viral copy number versus MFI values. The experiments were repeated five times for consistency.

### Combining microengraving and real-time PCR for clinical samples

We selected eight patients that originated from the Dominican Republic in the post-convalescent period of ZIKV infection. Demographic characteristics of analysed patients are shown in Table 1. ZIKV-infected patients were initially exanimated for developing a humoral response against ZIKV and presented high serological titer (IgG 8.54 - 20.4) using ELISA (Table 2). Further testing was performed to detect the presence of neutralizing antibody titer against ZIKV using PRNT. All patients had a neutralizing antibody titer, and six patients (45%) had a high score of neutralizing antibody titer against ZIKV (from 1:600 to 1:4000). All samples were negative for ZIKV using conventional real-time PCR. To detect and characterize antibody profiles in patients, we analyzed samples using the microengraving serological assay. As illustrated in Figure 3, capture slides coated with anti-human Ig/IgG (H+L) were hybridized with nanowell chips containing 10 μl of sample. After hybridization, the micrographs were processed for IgG, IgM, IgA antibodies and ZIKV envelope protein. All patient samples were positive for IgG, IgA, and ZIKV E protein and negative for IgM (Figure 2, Table 2). Since the samples were negative for ZIKV using conventional real-time PCR, we spiked the serum samples with ZIKV at different dilutions (1, 10, 100, and 1000 copies per nanowells predicted by Poisson distribution). The spiked samples were subjected to real-time PCR using the nanowells, and similar to Figure 2, the spiked samples were positive for ZIKV with a positive correlation (data not shown). These results demonstrate the ability to simultaneously detect ZIKV-specific antibodies and RNA using the nanowell chip assay.

**Figure 3.**
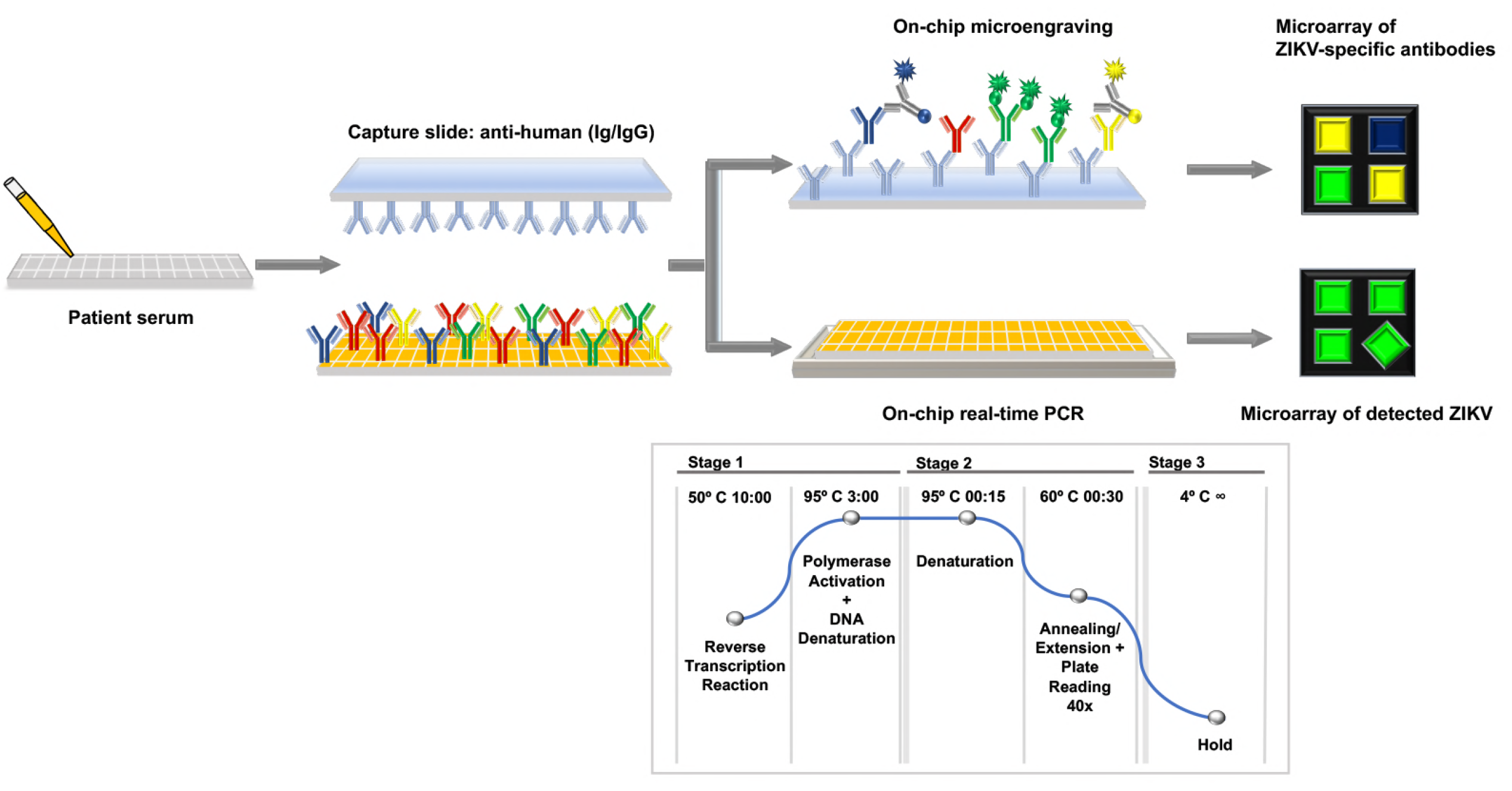
Schematic of testing patient sera using the combined on-chip microengraving and real-time PCR. Serum samples (n=8) were deposited into the nanowells. Microengraving was performed to detect anti-ZIKV E antibody, and IgG, IgM, and IgA isotypes. On the same nanowell chip, real-time PCR was conducted to quantify the viral loads. For this study, the serum samples were post-convalescent, therefore negative for ZIKV using conventional and on-chip real-time PCR. As a proof-of-concept, serum samples that were positive for anti-ZIKV E antibody (n=8) were spiked with different dilutions of ZIKV as presented in Figure 2 and on-chip real-time PCR was performed. Both experiments were repeated five times for consistency.

**Table 1.** Patients’ demographic profile

| Subject | Gender | Age (y) | Ethnicity | Origin Country | Suspected infection |
| --- | --- | --- | --- | --- | --- |
| 1 (plasma) | F | 35 | hispanic | Dominican Republic | ZIKV |
| 2 (plasma) | F | 54 | hispanic | Dominican Republic | ZIKV |
| 3 (plasma) | F | 23 | hispanic | Dominican Republic | ZIKV |
| 4 (plasma) | M | 62 | hispanic | Dominican Republic | ZIKV |
| 5 (plasma) | F | 47 | hispanic | Dominican Republic | ZIKV |
| 6 (serum) | F | 26 | hispanic | Dominican Republic | ZIKV |
| 7 (serum) | F | 35 | hispanic | Dominican Republic | ZIKV |
| 8 (serum) | F | 40 | hispanic | Dominican Republic | ZIKV |

**Abbreviations:** M= Male; F= Female.; y= years; ZIKV= Zika virus; DENV= Dengue virus.

**Abbreviations**: M= Male; F= Female.; y= years; ZIKV= Zika virus; DENV= Dengue virus.

**Table 2.**
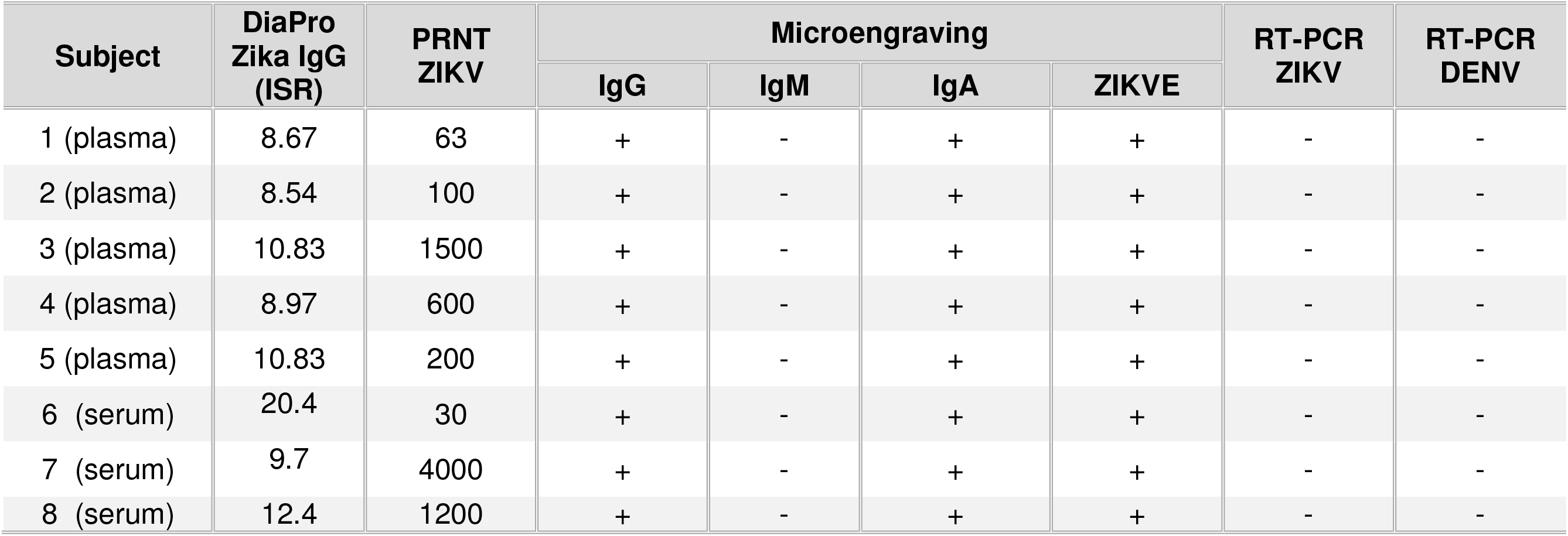
Patients laboratory diagnostic profile

**Abbreviations**: ISR= immune status ratio; IgG = immunoglobulin G; IgM = immunoglobulin M; IgA= immunoglobulin A; PRNT= plaque-reduction neutralization test; RT-PCR = realtime-polymerase chain reaction; ZIKV= Zika virus; DENV= Dengue virus; N/A= not applicable.

## DISCUSSION

There are currently 5 serological assays and 14 molecular assays for ZIKV detection with FDA emergency use authorization (EUA). The serological assays measure the IgM response against either E or NS1 protein for acute infection. The molecular assays amplify the E, NS1, NS3, or prM genes of the virus to quantify the presence of ZIKV using the real-time PCR platform (37). These diagnostic assays are essential for proper understanding of the transmission and clinical disease manifestations of Zika infection. However, the current serological and molecular tests are performed separately on different technical platforms which require larger sample volumes, is labor intensive, time consuming, and it carries a high degree of technical variability. In this study, to circumvent the inherent technical challenges, we developed an on-chip method for detection of ZIKV and anti-ZIKV antibodies with specific isotypes simultaneously using an array of nanowells. Our results demonstrate that by using microengraving as a serological test and real-time PCR as a molecular test, we were able to simultaneously detect isotypic antibodies against E protein and viral load of ZIKV at low copy numbers.

The ZIKV MAC-ELISA (CDC) serological assays use the recombinant, non-infectious ZIKV-like particles as capture antigen and demonstrated 94% positive agreement with PRNT and 83-100% positive agreement with peer-reviewed studies independently assessing the performance of the assays (37). Similar to the on-chip microengraving, the ZIKV Detect IgM ELISA (InBios, Seattle, WA, USA) utilizes the E protein as capture antigen. This assay showed 100% positive agreement with PRNT and 100% positive agreement with peer-reviewed studies (37). Using fluorochrome-conjugated ZIKV E protein as detection, our on-chip microengraving demonstrated significant specificity in which only gold standard anti-ZIKV E protein was detected, whereas negative control anti-flavivirus clone D1-4G2-4-15 performed as expected. To further test the on-chip microengraving, we analyzed plasma and serum samples from 8 post-convalescent patients (≥ one-year post infection). As demonstrated, these patients exhibited positive PRNT at various titers and due to the extended duration of time post-acute infection, these samples were all positive for IgG and IgA, but negative for IgM by standard ELISA. Similarly, our result demonstrates sera were positive for anti-ZIKV E, IgA and IgG, and negative for IgM.

The challenges of molecular assays to detect ZIKV RNA are that a low limit of detection can result in a high proportion of false negative results and testing conditions (samples, assay reagents, and experimental design) can compromise the sensitivity and specificity of detection. Available real-time PCR kits have varying limits of detection from 30 to 1000 copies/mL (38–40). The Trioplex RT-PCR (CDC) assay detects the ZIKV E gene using TaqMan real-time PCR with 100% positive agreement and a limit of detection of 1.93 × 10^4^ genome copy equivalents (GCE)/ml. Using the same primers and probes with modified reporters and quenchers, the on-chip assays were able to measure detectable fluorescent signals at one copy and showed positive correlation to 10^4^ copies. To examine the specificity of the test, DENV samples were also tested, and no signals were detected (data not shown). Therefore, real-time PCR on the chip in conjunction with epifluorescence microscopy exhibited remarkable sensitivity and specificity that are comparable or surpass current molecular assays in the market.

Due to the unfavorable consequences of ZIKV infection for populations living in areas endemic for ZIKV, and especially for pregnant women with a high risk of fetal abnormalities, rapid, highly specific, and sensitive diagnostic assays are urgently needed. The rapid outbreak and severe clinical manifestations launch a great urgency in the development of diagnostic tests for ZIKV. Zhang and colleagues proposed using a simultaneous serological assay on a nanostructured plasmonic gold platform for the detection of IgG, IgA, and IgG avidity against ZIKV and DENV-2 antigens in serum samples (41). Simultaneous detection of ZIKV, DENV, and Chikungunya based on the reverse transcription-loop mediated isothermal amplification (RT-LAMP) was proposed (42–44) with possible smartphone imaging (43). This one-step nucleic acid amplification method based on the PCR diagnostic test has many advantages such as rapidity of analysis and utilizing a portable format. Our on-chip detection method is limited by the high cost of instruments and technical expertise required. Part of our future study is to develop a portable on-chip process that is affordable and applicable in the field to monitor acute Zika infection.

## ACKNOWLEDGMENTS

The research was supported in part by Florida Department of Health, Biomedical Research Program (7ZK12, CQN) and the National Institute of Health (R21AI130561, CQN). The funders had no role in study design, data collection and interpretation, or the decision to submit the work for publication. The authors report no conflicts of interest. The authors have no competing financial interests regarding the subjects in this study

## REFERENCES

1. Cao-Lormeau VM, Roche C, Teissier A, Robin E, Berry AL, Mallet HP, Sall AA, Musso D. 2014. Zika virus, French polynesia, South pacific, 2013. Emerg Infect Dis 20:1085–6.

2. Mallet HP VA MD. 2015. Bilan de L’épidémie a virus Zika en Polynésie Française, 2013-2014. Bulletin D’linformation Sanitaires, Epidemiologiques et Statistiques 13:1–5.

3. CDC. 2016. All countries & territories with active Zika virus transmission https://www.cdc.gov/zika/geo/active-countries.html

4. Demir T, Kilic S. 2016. Zika virus: a new arboviral public health problem. Folia Microbiol (Praha) 61:523–527.

5. ECfDPa C. 2015. Rapid risk assessment. Zika virus epidemic in the Americas: potential association with microcephaly and Guillain-Barré syndrome.

6. Gatherer D, Kohl A. 2016. Zika virus: a previously slow pandemic spreads rapidly through the Americas. J Gen Virol 97:269–73.

7. Musso D, Nilles EJ, Cao-Lormeau VM. 2014. Rapid spread of emerging Zika virus in the Pacific area. Clin Microbiol Infect 20:595–6.

8. ECfDPa. C. 2016. Communicable disesases threat report week 6. http://ecdc.europa.eu/en/publications/Publications/communicable-disease-threats-report-13-feb-2016.pdf.

9. Martines RB, Bhatnagar J, Keating MK, Silva-Flannery L, Muehlenbachs A, Gary J, Goldsmith C, Hale G, Ritter J, Rollin D, Shieh WJ, Luz KG, Ramos AM, Davi HP, Kleber de Oliveria W, Lanciotti R, Lambert A, Zaki S. 2016. Notes from the Field: Evidence of Zika Virus Infection in Brain and Placental Tissues from Two Congenitally Infected Newborns and Two Fetal Losses–Brazil, 2015. MMWR Morb Mortal Wkly Rep 65:159-60.

10. Mlakar J, Korva M, Tul N, Popovic M, Poljsak-Prijatelj M, Mraz J, Kolenc M, Resman Rus K, Vesnaver Vipotnik T, Fabjan Vodusek V, Vizjak A, Pizem J, Petrovec M, Avsic Zupanc T. 2016. Zika Virus Associated with Microcephaly. N Engl J Med 374:951–8.

11. Oehler E, Watrin L, Larre P, Leparc-Goffart I, Lastere S, Valour F, Baudouin L, Mallet H, Musso D, Ghawche F. 2014. Zika virus infection complicated by Guillain-Barre syndrome–case report, French Polynesia, December 2013. Euro Surveill 19.

12. Kuno G, Chang GJ. 2007. Full-length sequencing and genomic characterization of Bagaza, Kedougou, and Zika viruses. Arch Virol 152:687–96.

13. Wang A, Thurmond S, Islas L, Hui K, Hai R. 2017. Zika virus genome biology and molecular pathogenesis. Emerg Microbes Infect 6:e13.

14. Kodati S, Palmore TN, Spellman FA, Cunningham D, Weistrop B, Sen HN. 2017. Bilateral posterior uveitis associated with Zika virus infection. Lancet 389:125–126.

15. Atkinson B, Thorburn F, Petridou C, Bailey D, Hewson R, Simpson AJ, Brooks TJ, Aarons EJ. 2017. Presence and Persistence of Zika Virus RNA in Semen, United Kingdom, 2016. Emerg Infect Dis 23:611–615.

16. Murray KO, Gorchakov R, Carlson AR, Berry R, Lai L, Natrajan M, Garcia MN, Correa A, Patel SM, Aagaard K, Mulligan MJ. 2017. Prolonged Detection of Zika Virus in Vaginal Secretions and Whole Blood. Emerg Infect Dis 23:99–101.

17. Calvet G, Aguiar RS, Melo ASO, Sampaio SA, de Filippis I, Fabri A, Araujo ESM, de Sequeira PC, de Mendonca MCL, de Oliveira L, Tschoeke DA, Schrago CG, Thompson FL, Brasil P, Dos Santos FB, Nogueira RMR, Tanuri A, de Filippis AMB. 2016. Detection and sequencing of Zika virus from amniotic fluid of fetuses with microcephaly in Brazil: a case study. Lancet Infect Dis 16:653–660.

18. Levi ME. 2017. Zika virus: a cause of concern in transplantation? Curr Opin Infect Dis 30:340–345.

19. Prevention UCfDCa. 2017. Guidance for US Laboratories Testing for Zika Virus Infection July 24, 2017.

20. Duffy MR, Chen TH, Hancock WT, Powers AM, Kool JL, Lanciotti RS, Pretrick M, Marfel M, Holzbauer S, Dubray C, Guillaumot L, Griggs A, Bel M, Lambert AJ, Laven J, Kosoy O, Panella A, Biggerstaff BJ, Fischer M, Hayes EB. 2009. Zika virus outbreak on Yap Island, Federated States of Micronesia. N Engl J Med 360:2536–43.

21. Schaub B, Vouga M, Najioullah F, Gueneret M, Monthieux A, Harte C, Muller F, Jolivet E, Adenet C, Dreux S, Leparc-Goffart I, Cesaire R, Volumenie JL, Baud D. 2017. Analysis of blood from Zika virus-infected fetuses: a prospective case series. Lancet Infect Dis 17:520–527.

22. Lanciotti RS, Kosoy OL, Laven JJ, Velez JO, Lambert AJ, Johnson AJ, Stanfield SM, Duffy MR. 2008. Genetic and serologic properties of Zika virus associated with an epidemic, Yap State, Micronesia, 2007. Emerg Infect Dis 14:1232–9.

23. Krow-Lucal ER, Biggerstaff BJ, Staples JE. 2017. Estimated Incubation Period for Zika Virus Disease. Emerg Infect Dis 23:841–845.

24. Baud D, Gubler DJ, Schaub B, Lanteri MC, Musso D. 2017. An update on Zika virus infection. Lancet 390:2099–2109.

25. Suy A, Sulleiro E, Rodó C, Vázquez É, Bocanegra C, Molina I, Esperalba J, Sánchez-Seco MP, Boix H, Pumarola T, Carreras E. 2016. Prolonged Zika Virus Viremia during Pregnancy. N Engl J Med 375:2611–2613.

26. Driggers RW, Ho CY, Korhonen EM, Kuivanen S, Jääskeläinen AJ, Smura T, Rosenberg A, Hill DA, DeBiasi RL, Vezina G, Timofeev J, Rodriguez FJ, Levanov L, Razak J, Iyengar P, Hennenfent A, Kennedy R, Lanciotti R, du Plessis A, Vapalahti O. 2016. Zika Virus Infection with Prolonged Maternal Viremia and Fetal Brain Abnormalities. N Engl J Med 374:2142–51.

27. Goncalves A, Peeling RW, Chu MC, Gubler DJ, de Silva AM, Harris E, Murtagh M, Chua A, Rodriguez W, Kelly C, Wilder-Smith A. 2017. Innovative and new approaches to laboratory diagnosis of Zika and dengue: a meeting report. J Infect Dis doi:10.1093/infdis/jix678.

28. Terzian ACB, Estofolete CF, Alves da Silva R, Vaz-Oliani DCM, Oliani AH, Brandão de Mattos CC, Carlos de Mattos L, Rahal P, Nogueira ML. 2017. Long-Term Viruria in Zika Virus-Infected Pregnant Women, Brazil, 2016. Emerg Infect Dis 23:1891–1893.

29. Gonzalez-Escobar G VA, Adams R, Polson-Edwards K, Hinds AQJ, Misir A, et al. 2017. Prolonged Zika virus viremia in a patient with Guillain-Barré syndrome in Trinidad and Tobago. Rev Panam Salud Publica.

30. Paz-Bailey G, Rosenberg ES, Doyle K, Munoz-Jordan J, Santiago GA, Klein L, Perez-Padilla J, Medina FA, Waterman SH, Gubern CG, Alvarado LI, Sharp TM. 2017. Persistence of Zika Virus in Body Fluids - Preliminary Report. N Engl J Med doi:10.1056/NEJMoa1613108.

31. Waggoner JJ, Gresh L, Mohamed-Hadley A, Ballesteros G, Davila MJ, Tellez Y, Sahoo MK, Balmaseda A, Harris E, Pinsky BA. 2016. Single-Reaction Multiplex Reverse Transcription PCR for Detection of Zika, Chikungunya, and Dengue Viruses. Emerg Infect Dis 22:1295–7.

32. Administration USFaD. 2012. 510(k) substantial equivalence determination decision summary, K113336, p 22, U.S. Food and Drug Administration, Washington, DC ed.

33. Langford MP, Weigent DA, Stanton GJ, Baron S. 1981. Virus plaque-reduction assay for interferon: microplaque and regular macroplaque reduction assays. Methods Enzymol 78:339–46.

34. Nguyen CQ, Ogunniyi AO, Karabiyik A, Love JC. 2013. Single-Cell Analysis Reveals Isotype-Specific Autoreactive B Cell Repertoires in Sjögren’s Syndrome. PLoS ONE 8:e58127.

35. Esfandiary L. VA, Ketchum J.M., Gupta N., Chan E.K., Stewart C.M., Bhattacharyya I., Sukumaran S., Nguyen C.Q. Development of a highly sensitive single-cell multiplex technology for early detection of Sjogren’s Syndrome, p. In (ed).

36. Love JC, Ronan JL, Grotenbreg GM, van der Veen AG, Ploegh HL. 2006. A microengraving method for rapid selection of single cells producing antigen-specific antibodies. Nat Biotechnol 24:703–7.

37. Theel ES, Hata DJ. 2018. Diagnostic Testing for Zika Virus: a Postoutbreak Update. J Clin Microbiol 56.

38. L’Huillier AG, Lombos E, Tang E, Perusini S, Eshaghi A, Nagra S, Frantz C, Olsha R, Kristjanson E, Dimitrova K, Safronetz D, Drebot M, Gubbay JB. 2017. Evaluation of Altona Diagnostics RealStar Zika Virus Reverse Transcription-PCR Test Kit for Zika Virus PCR Testing. J Clin Microbiol 55:1576–1584.

39. Frankel MB, Pandya K, Gersch J, Siddiqui S, Schneider GJ. 2017. Development of the Abbott RealTime ZIKA assay for the qualitative detection of Zika virus RNA from serum, plasma, urine, and whole blood specimens using the m2000 system. J Virol Methods 246:117–124.

40. Theel ES, Hata DJ. 2018. Diagnostic Testing for Zika Virus: A Post-Outbreak Update. J Clin Microbiol doi:10.1128/JCM.01972-17.

41. Zhang B, Pinsky BA, Ananta JS, Zhao S, Arulkumar S, Wan H, Sahoo MK, Abeynayake J, Waggoner JJ, Hopes C, Tang M, Dai H. 2017. Diagnosis of Zika virus infection on a nanotechnology platform. Nat Med 23:548–550.

42. Ganguli A, Ornob A, Yu H, Damhorst GL, Chen W, Sun F, Bhuiya A, Cunningham BT, Bashir R. 2017. Hands-free smartphone-based diagnostics for simultaneous detection of Zika, Chikungunya, and Dengue at point-of-care. Biomed Microdevices 19:73.

43. Priye A, Bird SW, Light YK, Ball CS, Negrete OA, Meagher RJ. 2017. A smartphone-based diagnostic platform for rapid detection of Zika, chikungunya, and dengue viruses. Sci Rep 7:44778.

44. Lamb LE, Bartolone SN, Tree MO, Conway MJ, Rossignol J, Smith CP, Chancellor MB. 2018. Rapid Detection of Zika Virus in Urine Samples and Infected Mosquitos by Reverse Transcription-Loop-Mediated Isothermal Amplification. Sci Rep 8:3803.

